# Sinogram Domain Angular Upsampling of Sparse-View Micro-CT with Dense Residual Hierarchical Transformer and Noise-Aware Loss

**DOI:** 10.1101/2023.05.09.540072

**Authors:** Amogh Subbakrishna Adishesha, Daniel J Vanselow, Patrick La Riviere, Keith C Cheng, Sharon X Huang

## Abstract

Reduced angular sampling is a key strategy for increasing scanning efficiency of micron-scale computed tomography (micro-CT). Despite boosting throughput, this strategy introduces noise and artifacts due to undersampling. In this work, we present a solution to this issue, by proposing a novel Dense Residual Hierarchical Transformer (**DRHT**) network to recover high-quality sinograms from 2 ×, 4× and 8× undersampled scans. DRHT is trained to utilize limited information available from sparsely angular sampled scans and once trained, it can be applied to recover higher-resolution sinograms from shorter scan sessions. Our proposed DRHT model aggregates the benefits of a hierarchical-multi-scale structure along with the combination of local and global feature extraction through dense residual convolutional blocks and non-overlapping window transformer blocks respectively. We also propose a novel noise-aware loss function named **KL-L1** to improve sinogram restoration to full resolution. KL-L1, a weighted combination of pixel-level and distribution-level cost functions, leverages inconsistencies in noise distribution and uses learnable spatial weights to improve the training of the DRHT model. We present ablation studies and evaluations of our method against other state-of-the-art (SOTA) models over multiple datasets. Our proposed DRHT network achieves an average increase in peak signal to noise ratio (PSNR) of 17.73dB and a structural similarity index (SSIM) of 0.161, for 8× upsampling, across the three unique datasets, compared to their respective Bicubic interpolated versions. This novel approach can be utilized to decrease radiation exposure to patients and reduce imaging time for large-scale CT imaging projects.

## 1 Introduction

Synchrotron micro-CT optimized for whole organisms at sub-cellular resolution (histotomography) has been proposed as the foundational tool for computational tissue phenotyping [7]. The large datasets generated by this approach are well suited for analysis with modern computational techniques such as cell detection and shape estimation, and they could prove useful for registration of multi-modal data, which can then lead to significant biological insights [19]. However, the acquisition process involved can be a hindrance to research that requires high throughput, as imaging large specimens may require significant ‘beam time’ for optimal scan quality. We wish to alleviate this bottleneck and enable faster scanning without compromising scan quality [9].

At synchrotron sources, it is most common for the serial rotational 2D projections to be taken as the subject is continuously rotated. These projections are then processed using filtered backprojection (FBP) to obtain a digital 3D reconstruction of the sample. This setup is illustrated in **Figure 1**. Long scans, though vital for acquiring high-resolution sub-cellular details, limit the number of samples that can be scanned during a typical beamtime allocation. Long scans also require sophisticated equipment for stabilizing and monitoring the sample during imaging. Non-orthogonal and inconsistent positioning leads to significant image artifacts, particularly in our setting of large-field, high-resolution micro CT [34].

**Figure 1.**
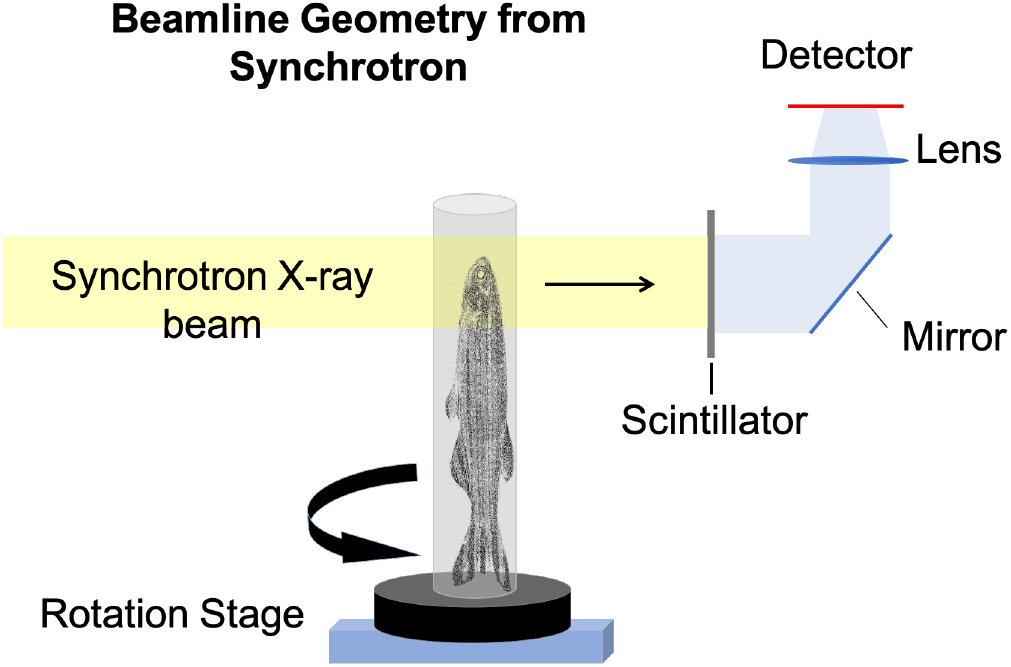
Parallel beam Micro-CT image acquisition by rotation of subject

Two immediate approaches for decreasing the scan times are available: **a)** Reducing the X-ray exposure time at each of these angles and denoising the resulting scans as demonstrated in [38],[41] and [6] or **b)** Reducing the number of sampling angles around the subject as performed in [11] and [27]. In the former technique, since fewer X-ray photons reach the sensor, there is an increase of Poisson-based noise in the scans that can severely deteriorate the quality of the scans and can obscure biologically relevant sub-cellular details. The latter too presents challenges of its own. Due to the sparsity in the information available to the FBP algorithm, several types of artifacts are introduced in the reconstructed volumes: stretching, streaking, and rotational blur, all of which adversely affect their readability.

In this work, we aim to address the concern of under-sampling artifacts and noise of approach **b)** through an innovative deep-learning network. We propose a novel Dense Residual Hierarchical Transformer (DRHT) to interpolate the information available in the angular axis to recover a high-resolution scan from an under sampled (reduced angle) scan. The dense and residual structures of the DRHT network successfully capture local feature interactions while the non-overlapping window-based attention blocks acquire the global feature interactions across multiple scales of the hierarchical U-shaped architecture. We present detailed ablation studies to show the need for each of the constituent sub-units in our network. In addition to the various novelties presented by our network pipeline, we propose a novel noise-aware learned-weighted loss combination named *KL* − *L*1 (where KL is the Kullback–Leibler divergence and L1 is the Mean Absolute Error) loss to overcome the challenges posed by non-uniform information distribution in under-sampled sinograms. Our DRHT along with *KL*− *L*1 loss empirically outperforms existing state of the art models aimed at performing single-image sinogram angular upsampling for sparse-view micro-CT. In the spirit of reproducibility and transparency, we shall make our code publicly available on Github. Our contributions can be highlighted as follows:

- We present a novel sparse-view CT image restoration method in the sinogram domain which can accurately restore images to full angular resolution from 2×, 4× and 8× under-sampled versions.
- To perform accurate sinogram restoration, we propose a novel dense-residual hierarchical transformer (DRHT) network built with highly functional and modular sub-units which effectively removes artifacts and improves signal to noise ratio significantly.
- To improve the training, we present an advanced sinogram specific, noise-aware, KL-L1 weighted loss function capable of addressing areas with and without subject uniquely.
- Through a detailed multi-scale evaluation against existing models with both quantitative and qualitative measures we show superiority of the proposed DRHT model.

## 2 Related Work

Image super resolution or upsampling has received significant attention from both computer vision and biomedical imaging research communities. In the computer vision literature, among the initial works to use convolution neural networks (CNNs) for super resolution was [10] which presented the superiority of CNNs over the sparse-coding methods prevalent before them. Kim et al. [22] extended this to show that deeper networks like the VGG-Net performed better as they could capture finer features more efficiently. Following the proposal of Generative Adversarial Networks(GAN) in [13], many GAN based super resolution models were proposed. Among them, a popular work was by [24] which had an encoder-decoder network as a generator and a simple CNN as a discriminator. GANs gained popularity due to the robustness of the loss function and they even showed good success with reference based super resolution applications. One such notable work is [55] which uses a texture transfer technique between consecutive generators at multiple scales.

Deep residual neural networks, though originally proposed for image recognition in the work [17], proved excellent in propagating features from fine-grain details across multiple layers, primarily due to their skip connections, a vital requirement for super resolution. Residual Dense Network (RDN) [53] uses multiple stacked Residual Dense Blocks (RDBs) to perform single-image super resolution. If the image is resized to the target size before super resolution, the task becomes similar to noise removal as the network has only to learn the mapping parameters between the two similar-sized images. [52] performs this operation while using the same RDN for mapping. Video frame interpolation or temporal infilling using RDNs was done by [40] and [23].

Recently, transformers have been extensively used in low-level vision tasks like super resolution and denoising as notably presented by [4], [12], [29], [32] and [49]. Their ability to extract long-range feature interactions and use them in noise-free reconstruction has proven extremely successful for a wide variety of tasks. The Uformer proposed in [45] further improves the transformer’s abilities through a multi-scale hierarchical feature extraction and skip connections at similar resolutions to maintain sharpness while decoding. The skeleton of the architecture is similar to the original U-Nets proposed by [37]. Diffusion based models have been used in the field of super resolution [28] and [36]. However, current diffusion models are designed for uniform Gaussian noise priors while sub-sampled micro-CT have non-uniform Poisson priors with artifacts. For to this reason, we contend that diffusion based models are currently not applicable for our purpose.

Exploring the medical imaging domain, CT super resolution has been performed with CNN based networks in [35], [50] and [51] and with U-Net and sub-pixel based approaches in [16]. An interesting amalgamation of successful sub-components includes [14] which combines residual dense structures from RDNs with hierarchical units of U-Nets to achieve better image denoising results. To bolster the argument for U-Nets, [1] show that using a hierarchical structure yielded higher performance over their previous work [2] called ‘SRCN’. Chao *et al*. [42] used a Cycle-GAN (SRCGAN) technique to perform sinogram super resolution though at a risk of GAN related artifacts. While these works describe general sinogram-domain super resolution, there are works specifically in the context of sparse view that are more relevant to our contribution. For example, [54], [48] and [25] have proposed variations of deep neural networks to perform artifact and noise removal of sparse view CT in the reconstructed image domain. More recently, [46] and [18] have presented the two-model concept where the sinogram domain and the image domain are processed by separate networks, which increases the computational resources required to super-resolve the signal. We wish to solve this purely in the sinogram domain to ensure usability by researchers who may not have computational resources for using two deep learning models. Based on the review of the literature, a combination of vital feature-extracting elements such as dense residual structures for local feature extractions and non-overlapping window-based attention mechanisms for long-range interactions along with a hierarchical skeletal structure have the potential to achieve high-performance in the image restoration task. The novel combination proposed by our model thus ensures an improved single-image angular upsampling for sinogram-domain images.

In our work we are comparing the proposed novel DRHT against a Bicubic [21] baseline and recent state of the art single-step networks including RDN [53], RDUNET [14], REDCNN [5] and Uformer [45] models.

## 3 Methodology

Traditionally, super resolution is performed on 2D spatial row-column axes and most algorithms in the literature are designed around such problem statements. We convert our sub-sampled angular scans to spatial data using a sinogram-domain representation. A sinogram represents a complete set of 1D angular X-ray projections for a specified slice of an object. These 1D projections are then stacked in the row dimension to form a 2D sinogram. For example, given a set of 1500 2D projection images acquired over 180 degrees, we would extract the *n*th row from each such image and assemble these into a new 1500-row array representing the sinogram of slice *n* of the object. We can use this as our high resolution (HR) reference sinogram. Instead of re-acquiring reduced-angle scans with larger angular steps, we can drop alternate rows in the reference sinogram to obtain the 2× downsampled sinograms. We can continue dropping alternate rows to obtain subsequent 4× and 8× down-sampled sinograms. It is, however, important to note that only the rows are being halved and the columns do not change. This causes a 1500 ×2048 reference image to become 750× 2048, 375 ×2048 and so on. In order to recover the full-resolution reference, we need to map a rectangular region in the down sampled scan to a square region in the reference scan. A 64× 128 patch in the 2× downsampled sinogram maps to a 128 ×128 patch in the reference full sized sinogram. A simple workaround is to resize the downsampled sinograms to the same size as the reference and then use a 1:1 mapping between them. We use Bicubic interpolation to stretch the downsampled sinograms in the rows back to full resolution. This causes some stretching artifacts in the input sinograms and can be seen in the fourth column of **Figure 2**. Unlike traditional super-resolution where the noise and artifacts are uniformly distributed across the height and width of the image, in sinograms, due to angular sampling, the inconsistencies and sparsity increase radially further away from the centre of object’s axis of rotation as illustrated in the first column of **Figure 2**. We address this in detail with our proposed approach in Section 3.3.

**Figure 2.**
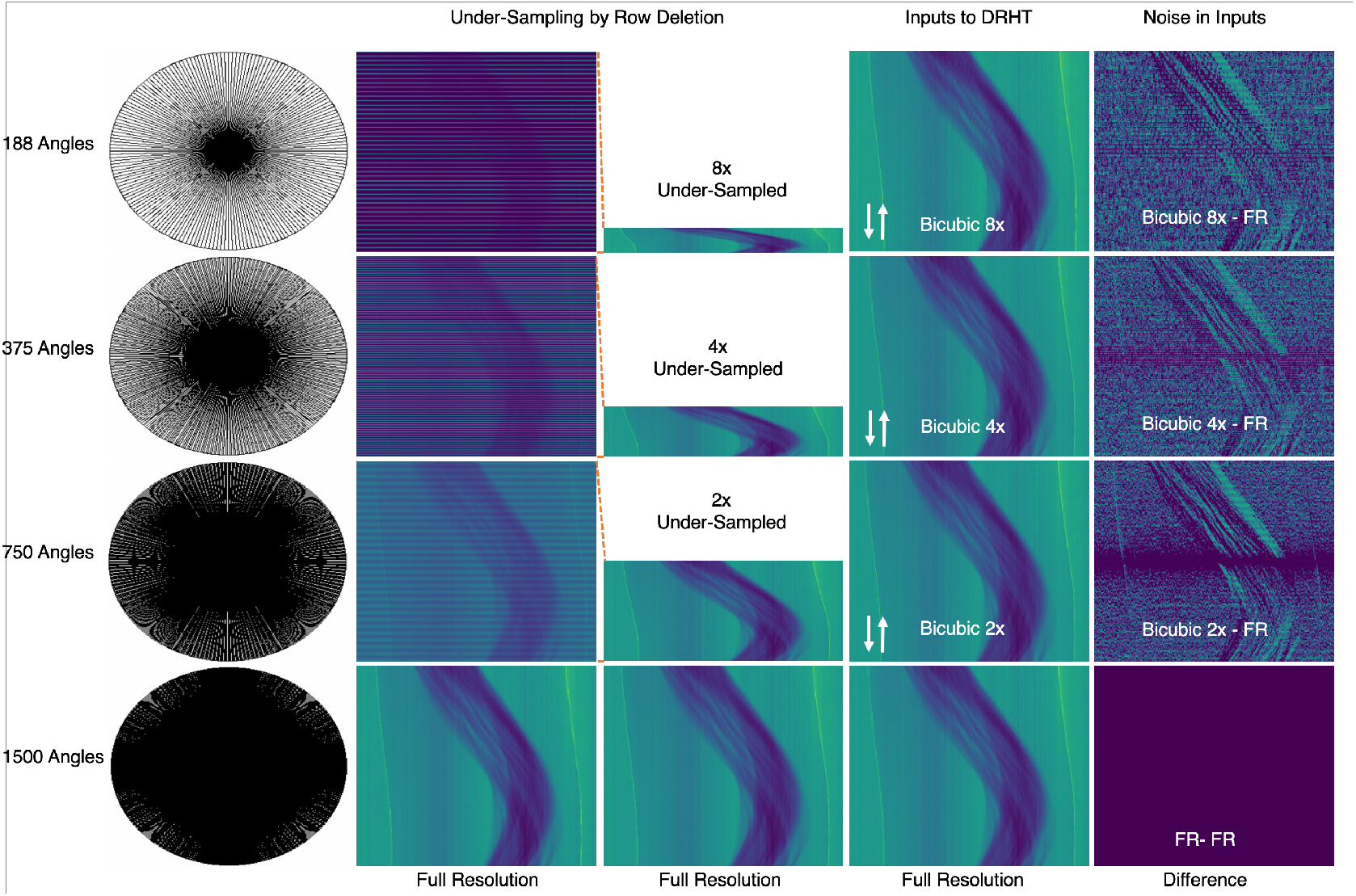
First column illustrates the angular sampling pattern of a subject over 1500 (full), 750 (half), 375 (quarter) and 188 (eighth) angles. Note the presence of darker region (over-sampled) and the sparse region (under-sampled) in the circles. The second and third columns show the under-sampled signal from the Zebrafish dataset obtained by row deletions in the sinogram. The fourth column is the input to the DRHT network where the original size is re-obtained using Bicubic interpolation algorithm. Here ⇵ indicates the interpolation operation on the under-sampled signal. The last column illustrates the difference between the DRHT inputs at different scales and the full-resolution ground truth.

### 3.1 Data

We utilize three publicly available and published datasets where the raw 2D projections and the reconstruction parameters are made available. We transpose the projections into a set of 2D sinograms for each volume. The first dataset is a cone-beam X-ray CT scan, [8], of a walnut acquired at 100*µ*m resolution with 1201 projections across the angular space. A total of 42 walnuts are available on Zenodo^1^ of which we use 8 walnuts for training and test on 2 held-out walnuts using 5-fold cross validation. Our second dataset is an Earthworm [26] acquired at 8.17*µ*m resolution with 960 projection images. The data can be found at GigaDB^2^. In this dataset, we extracted 35840 sinogram patches for training and 8960 non-overlapping patches from the same set for testing. Finally, we include a dataset of Zebrafish larvae obtained 5 days post fertilization [9] acquired at 0.74*µ*m resolution with 1501 projections across the whole organism. The data is publicly available on Dryad^3^. Here, among the 5 fish sets available, we use 4 for training and reserve the held-out one for testing in a 5-fold cross validation method.

We first convert the projections to sinograms and then downsample them using row deletion described in the previous subsection. We repeat the alternate row deletion method to further degrade the image in order to simulate 4× and 8× undersampling. We then resize the down-sampled sinograms using Bicubic interpolation [21] back to the original size. Following this, we randomly sample 8 to 16 square patches of size 128× 128 from each sinogram. The numbers of patches used from each dataset for training and testing purposes are detailed in **Table 1**. While these downsampled-upsampled patches act as input, the original reference, which is unaltered at the corresponding location acts as the target patch.

**Table 1:** Micro-CT Datasets for Model Training and Evaluation

| Name | Specimen | Resolution | # of Projection Angles | # of Training Patches | # of Test Patches |
| --- | --- | --- | --- | --- | --- |
| Walnut Micro-CT [8] | Juglans regia | $100\mu\text{m}$ | 1201 | 24576 | 6144 |
| Earthworm Micro-CT [26] | Aporrectodea trapezoides | $8.17\mu\text{m}$ | 960 | 35840 | 8960 |
| Zebrafish Micro-CT [9] | Danio rerio | $0.74\mu\text{m}$ | 1501 | 65536 | 16384 |

### 3.2 Dense Residual Hierarchical Transformer Network (DRHT)

Our proposed DRHT model uses the Bicubic interpolated patches as input and generates clean, artifact-free patches of the same size as output. We train individual models for scale (2×, 4× and 8×) and test them on held-out data and report the performance on the test set. We use peak signal to noise ratio (PSNR) and structural similarity (SSIM) to compare the model performance. Our goal is to increase image quality with complete fidelity, that is, upsampling while taking great care not to introduce morphological features that do not exist in the sample.

The DRHT network, illustrated in **Figure 3**, can be described as a symmetric hierarchical structure with DRHT blocks at each scale, one in the encoding direction and another in the decoding direction. We use an additional DRHT block at the bottle-neck level of the network. In the following subsections, we detail the structure of the network and the sub-components of the DRHT block.

**Figure 3.**
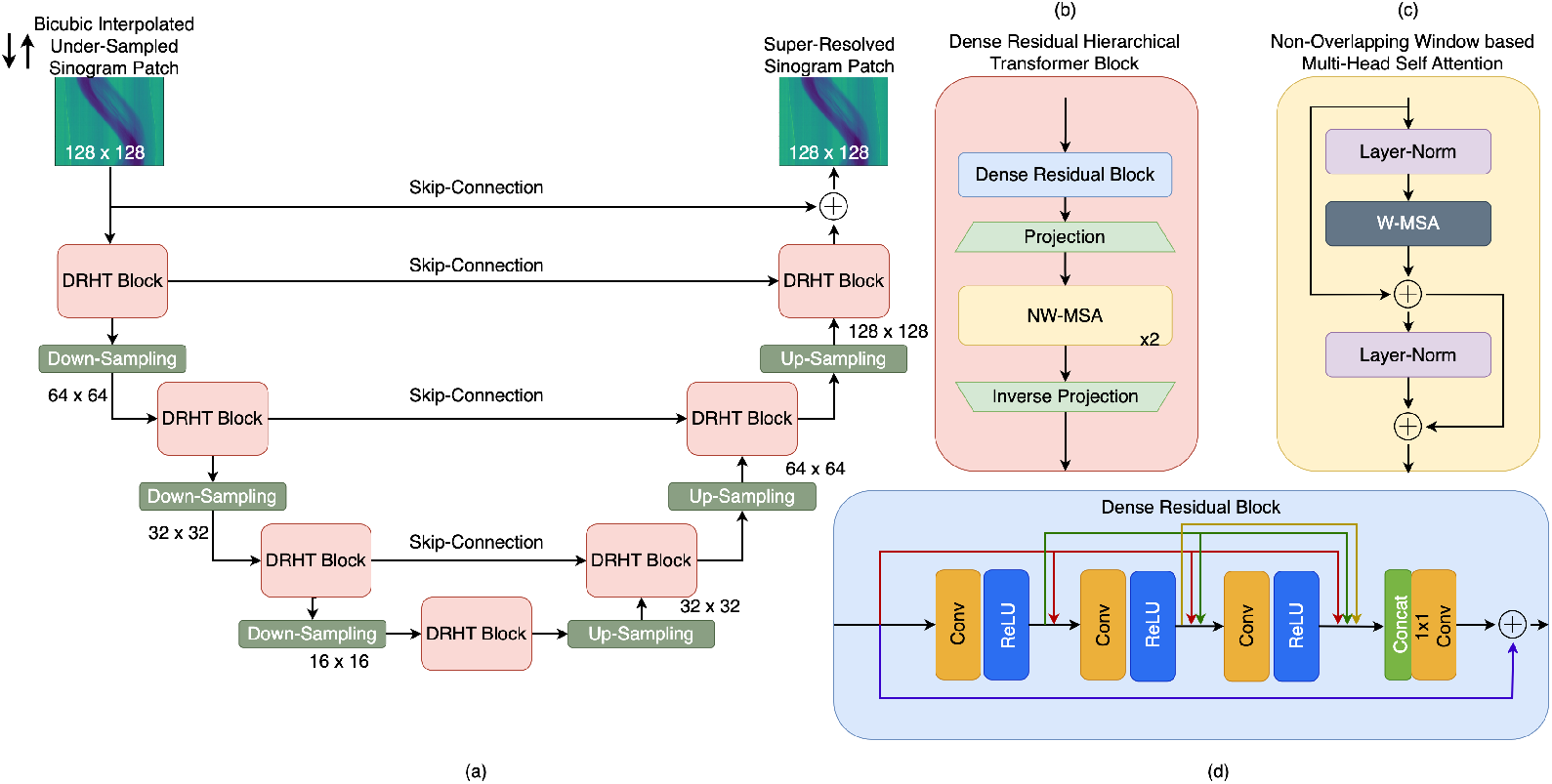
(a) Architecture of proposed Dense Residual Hierarchical Transformer (DRHT) Network; (b) DRHT block with a Dense Residual Block, Projection layers and Non-Overlapping Window based MSA (NW-MSA) blocks; (c) A unit of NW-MSA; (d) Dense Residual Block with three Conv-ReLU layers and a feature aggregation layer.

#### 3.2.1 Hierarchical Structure

First, the skeletal architecture of DRHT involves a U-shaped multi-scale structure with symmetric downsampling and upsampling operations. At each scale, a DRHT block is used to capture features while skip connections between symmetrically opposite encoder and decoders help improve sharpness for reconstruction. At the bottle-neck, we apply a similar DRHT block to process the signal at the lowest scale and capture global dependencies. The down-sampling is done using a 4 ×4 convolution kernel with stride set to 2, which halves the image in both height and width dimensions. Similarly, the upsampling is performed using a 2× upsampling layer followed by a convolution layer with a kernel size of 3 to avoid checkerboard artifacts. The output of the final decoder DRHT block is added to the initial input and the upsampled artifact-free image is produced.

#### 3.1.2 DRHT Block

A DRHT block comprises five components: a dense residual block (DRB), two projection layers, and two non-overlapping window-based multihead self-attention (NW-MSA) transformer blocks. The DRB is detailed in the following subsection. The first projection layer flattens the features for the transformer blocks while the inverse projection layer reshapes the features in order to perform the upsampling/down-sampling operations. Provided with an input of dimensions 1× H ×W, the projection layer, which has a convolutional kernel of size 3× 3, results in a set of shallow features of size C ×H ×W, where C is the number of channels for that scale. At each scale of the encoder, the number of channels is doubled while the height and width are halved. At the s-th stage, the data takes the shape of 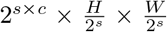 The decoder on the other hand performs halving of the channels while doubling the height and width dimensions.

#### 3.2.3 Dense Residual Block (DRB)

The DRB is a vital component of the DRHT comprising three stacked Conv-ReLu layers with both dense and residual connections across them. To avoid propagation of a large number of features, we aggregate them using a 1× 1 Conv layer that is then combined with the input feature vector. The DRB is responsible for both extracting local feature interactions and for propagating deep features. Deep neural networks are vulnerable to vanishing gradients and using DRBs helps avoid this and improves the overall training process.

#### 3.2.4 NW-MSA

In each of our transformer blocks, we use non-overlapping window-based multi-head self-attention layers positioned between two-layer norm blocks. There are residual connections to improve the flow of features through the transformer block. The NW-MSA has a predefined window size, *r*^2^, and image size, *hw*, at each level. The ratio of 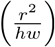 grows with each encoding level and decreases with each decoding level. At the initial levels, the transformer benefits from calculating attention only within a window, which helps extract local interactions, while closer to the bottle-neck level, the attention computed is nearly global as the size of the image is close to the window size. If each window provides a flattened and transposed feature vector *X* and we use a total of *k* attention heads, we can formulate the MSA output as follows:

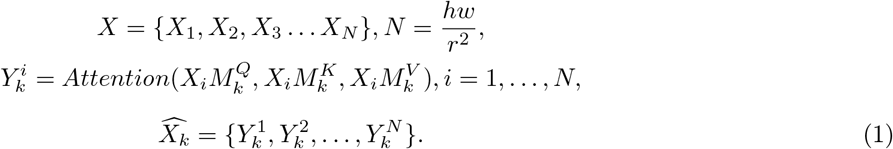

Here, 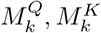 and 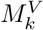 are the query (Q), key (K), and value (V) projection matrices for the *k*-th head. Outputs for each of the *k* heads are concatenated for the final output. This closely follows the window-based attention calculation established by [45]. To calculate the attention function, we refer to [30] and [39] where if 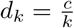 is the head dimension of the *k*-th head, then

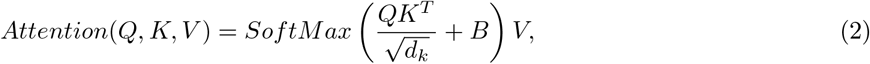

where *B* is the learnable relative positional bias term. We leverage the ability of the transformer block to recognize feature interactions at multiple scales to reconstruct clean upsampled sinograms. The DRHT model is trained for 100 epochs and the epoch with the best validation performance is picked for testing. AdamW [31] optimizer is used with an initial learning rate set to 0.0002 and gradually reduced using a step-wise decay function. The training is performed using 4 Nvidia Quadro RTX6000 GPUs.

### 3.3 Loss

Conventionally, for image restoration tasks, pixel-wise intensity based loss terms like mean absolute error (MAE), also known as *L*1, or mean squared error (MSE), also known as *L*2, are used to reduce the difference between the predicted and target images. This works well for natural images with uniform information distribution across the height and width of the image. In under-sampled sinogram images, a variety of factors additionally affect the noise profile. (1) The noise is signal dependent (Poisson based) and not uniform spatially. The standard deviation of the noise present is proportional to 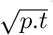where *p* is the expected number of photons per unit time at the specific detector and *t* is the exposure time. (2) Due to the angular sampling process, regions closer to the axis of rotation are sampled far more than the peripheral regions. Refer to **Figure 2** to observe this phenomenon. (3) L1 and MSE both average over the entire image. In our training, if a particular patch contains minimal subject information and is largely flat, this average tends to be very low. This low loss provides weaker updates to the network, which slows down the search for global minima and hence adversely affects the training process. (4) Traditionally, during training, the model learns to map each pixel intensity to another. However, in our setting, sinogram patches with largely flat areas (air/plastic) cannot be mapped to fixed intensities due to the presence of random noise. Instead it would be ideal to map them to a noise-free distribution. Our motivation to propose a weighted-learned loss function comes from these previously unaddressed issues.

We emphasize that due to the spatial variability of the noise profile, the PSNR also changes based on distance from the axis of rotation. **Figure 4** illustrates the peak signal-to-noise ratio (PSNR) distribution over the image. It can be observed that regions sampled closer to axis of rotation (with Zebrafish) have significantly higher PSNR compared to regions sampled further away from the center.

**Figure 4.**
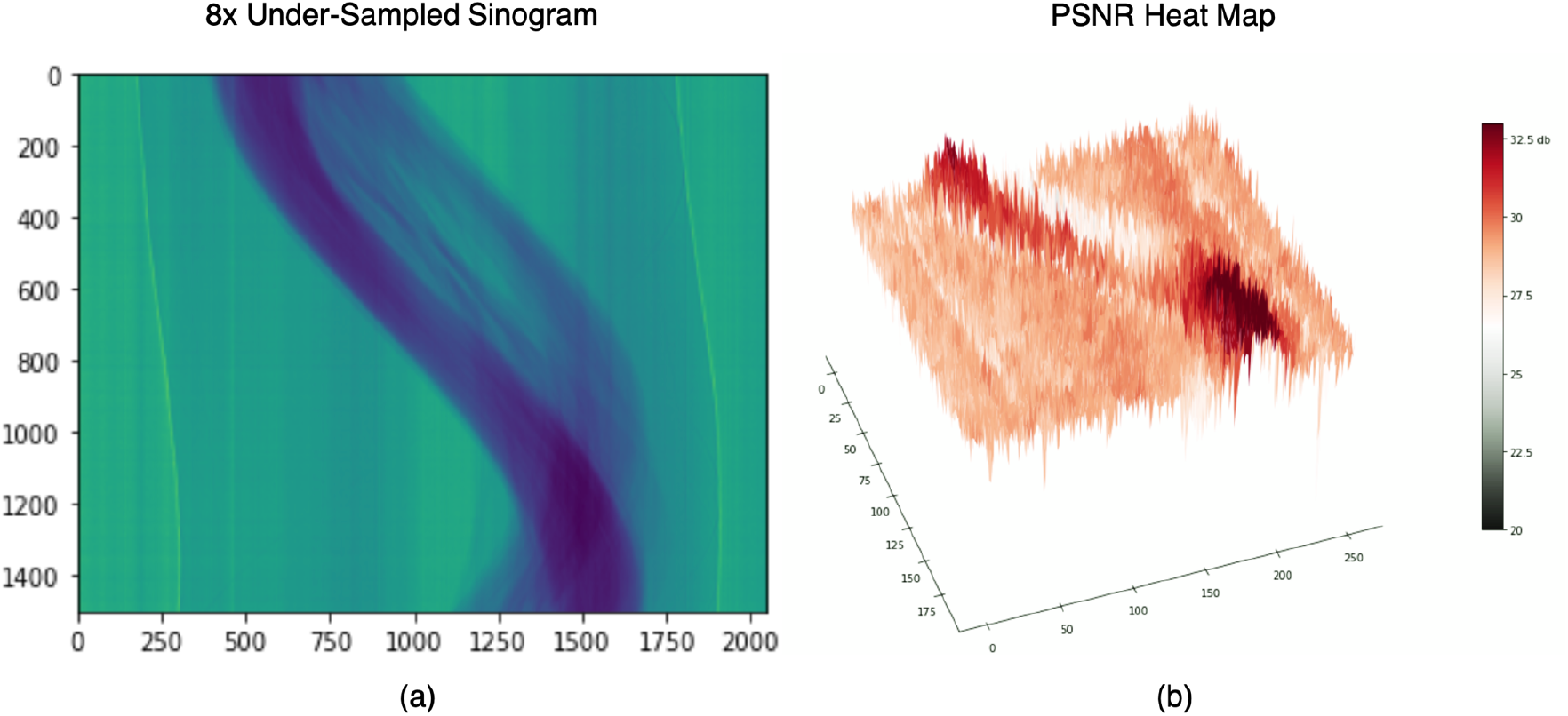
Motivation for a novel noise-aware loss (a) ⇵ Bicubic 8x under-sampled sinogram input from Zebrafish dataset; (b) PSNR of (a) with respect to full resolution reference calculated over blocks of 8×8 pixels. The regions with the subject has a higher PSNR than regions without the subject.

The above mentioned issues warrant a need for noise-aware loss functions that can pay attentions to specific areas within the image instead of the entire image. Since the spatial location of noise changes for each sinogram, a simple Gaussian filter like the one used by [3] will not suffice. To address the above mentioned issues, we propose a weighted KL-L1 loss which permits learning a higher L1 weight in regions with high details like the subject and a higher KL weight for flatter regions without (like plastic and background). We learn these weights during the training process and apply them individually for each input.

#### 3.3.1 KL-L1 Loss Intuition

As mentioned in previous subsection, the amount of information in the sinogram depends on the region in question. For example, flatter regions (plastic/air) do not contain any details and should ideally be of fixed intensity. However, due to randomness of noise and the presence of stretching artifacts, they contain local variations of intensity. While learning to accurately recover these flat regions, it is futile to expect each pixel to match that of ground truth. It instead is easier to learn the intensity distribution of the flat region and recreate the same in the restored image. For regions in the sinogram with subject, where every pixel is important, a stronger pixel-wise loss like L1 can help learn the accurate intensities. The resulting combination of the two can then leverage complementary benefits and in turn result in a cleaner and more uniform sinogram. The intuition behind the loss function is illustrated in **Figure 5**.

**Figure 5.**
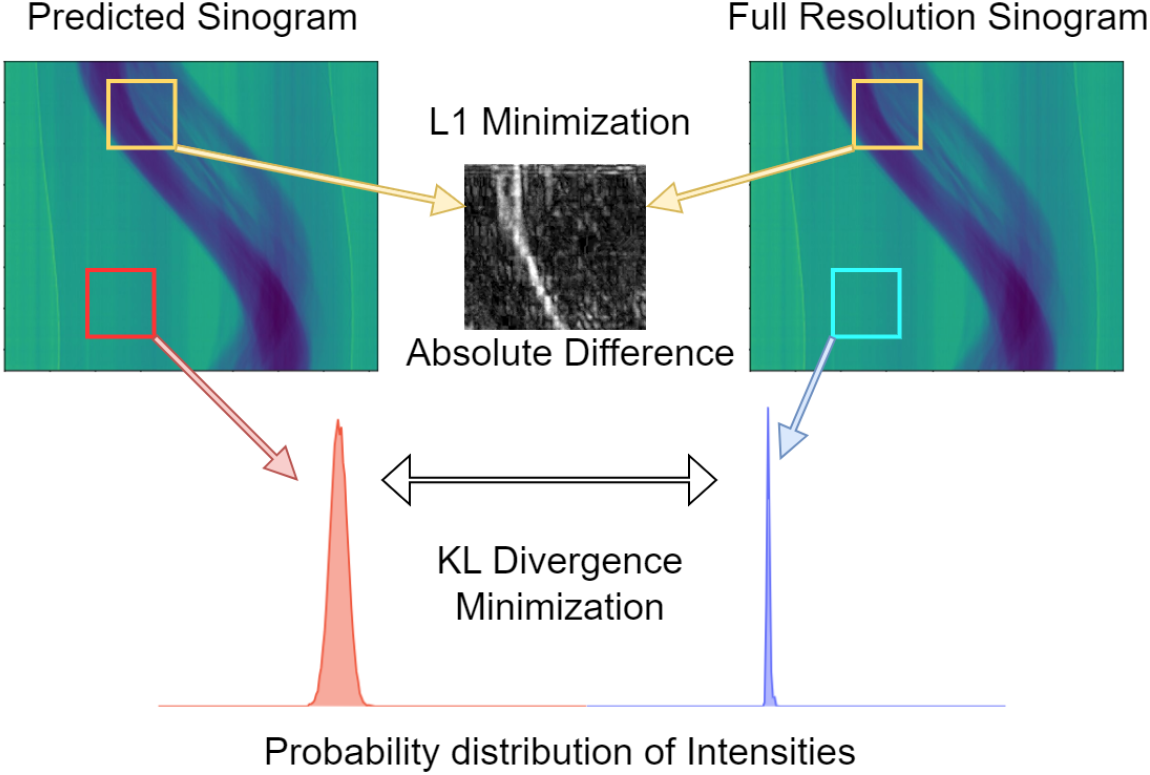
Different regions in the predicted sinogram can be learned using either distribution distance (KL) or pixel-level difference (L1) depending upon the information contained in them.

#### 3.3.2 Novel Weighted KL-L1 Loss Calculation

The loss calculation involves three sub-steps. (1) Calculation of a weighted L1-Loss and (2) Calculation of a weighted KL divergence loss and (3) The scaled averaging of the two. **Figure 6** illustrates this process in detail. In the first part, the model produces a 128× 128 weight matrix corresponding to the 128 ×128 sized input. A pixel-level absolute error is extracted and then an element-wise multiplication with the weight matrix is performed. If the pixel-level absolute error is termed *P*_*AE*_ and the L1 weight matrix is *W*_*L*1_, the element-wise product is (*P*_*AE*_.*W*_*L*1_). To prevent the loss from going to 0, we use a stabilization term 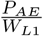 which is added to the previous result. Following this, the mean is calculated across the resulting matrix.

**Figure 6.**
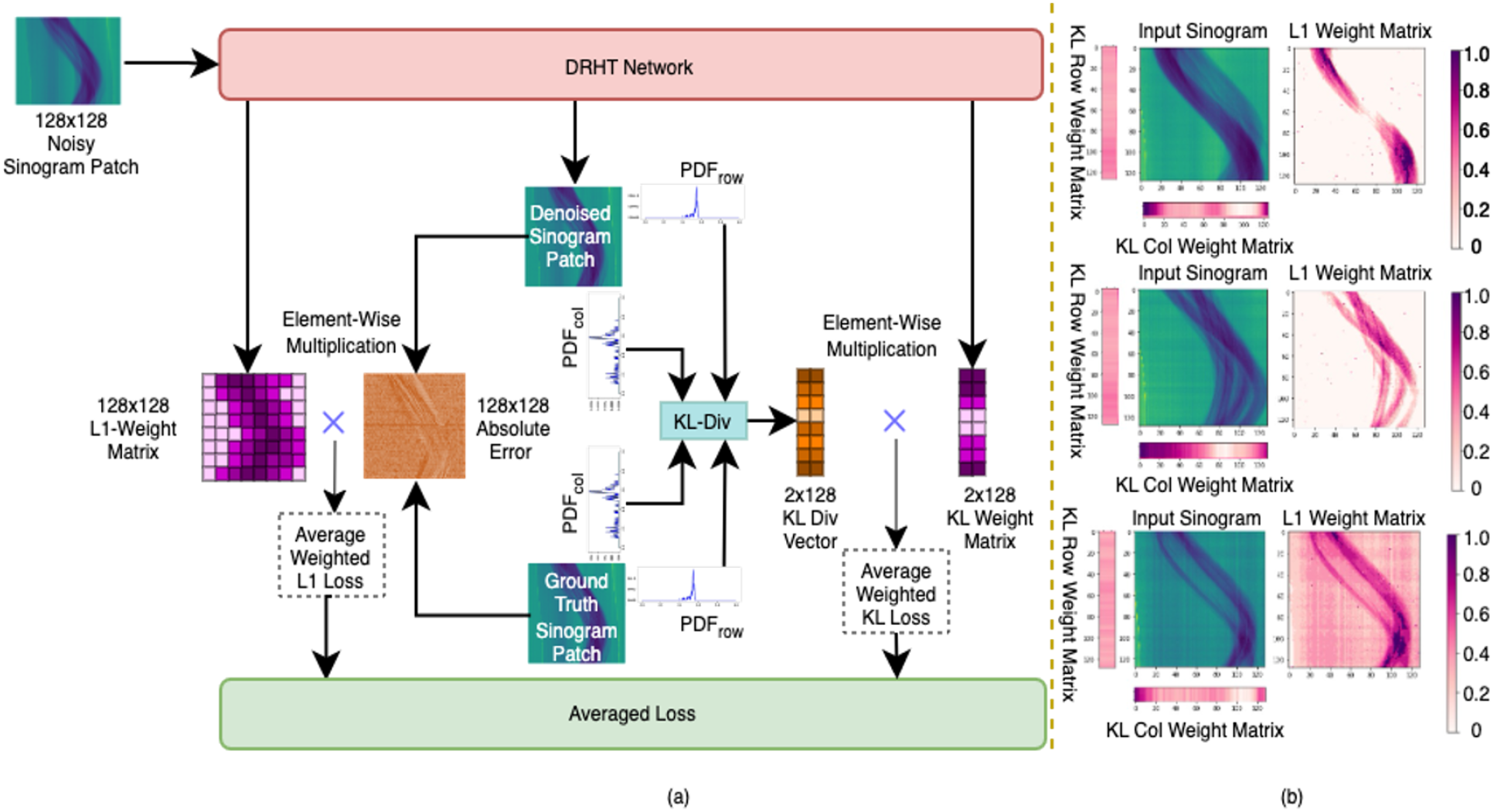
(a)Calculation of Weighted KL-L1 Loss. The absolute error across each pixel is multiplied with the L1 weight matrix of the same size. Similarly, row-wise and column-wise KL divergences are multiplied with their corresponding weight matrices. (b)Examples of Sinogram inputs and their respective L1 and KL weight matrices.

In the second part, KL divergence is an estimation of the distance between two probability distributions. An image can be reduced to a probability distribution through a SoftMax layer of an encoder network as performed in [47],[56]. However, calculating KL divergence between two entire images increases computational requirements. Additionally, the granularity of details present in local regions cannot be completely utilized when using the whole image instead of rich row-level and column-level data. Each column in the sinogram corresponds to sampling the subject at a fixed distance from multiple angles while each row corresponds to sampling different points of the subject from a fixed angle. To accommodate this specification, we compute row-wise and column-wise 100-binned histograms (using pre-determined minimum and maximum values) with a triangular kernel density estimation prescribed in [33]. Using the histograms, we extract the probability density functions (PDFs) for each row and each column from both the predicted image and the ground truth reference. If the size of the image is *M× N*, this operations results in *M* PDFs for the rows and *N* PDFs for the columns. If a row from the denoised image and the same row from the ground-truth image are considered, they ideally should have the same PDF. This is true for columns as well. However, due to the randomness of the noise present, the distribution varies. Considering a row *i* at a time, we calculate the KL-divergence between the PDF of row *i* from the predicted image and the PDF of row *i* from the reference image. We repeat this for all *M* rows and similarly extract KL divergences for each of the *N* column pair resulting in a 1×M sized vector for the rows and a 1 ×N sized vector for the columns. The DRHT network also produces a similar sized KL-weight matrix. Since we use 128 ×128 sized patches, the KL-weight matrix is 2× 128 on which element-wise multiplication and stabilization is performed before averaging. The two averages are then scaled with learned scales (*λ*_1_ and *λ*_2_) and averaged to form the final loss. From our experiments, the L1 weight matrix has higher weights from regions with subject and lower weights elsewhere. The KL-column weight matrix favors columns without subject while the KL-row weight matrix has no decipherable pattern. This is shown in **Figure 6b**. The weighted L1 loss is given by

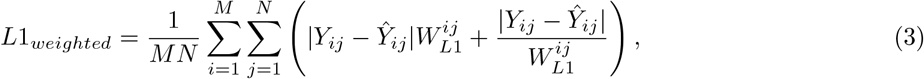

where *W*_*L*1_ is the *L*1 weight matrix, *M* is the number of rows and *N* is the number of columns and |*Y*_*ij*_− *Ŷ*_*ij*_| is the *P*_*AE*_ detailed earlier.

Following [43], we estimate histograms for each row and each column in the image using a triangular kernel density function. If 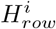 is the 100-binned histogram of row *i* and 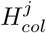 is the 100-binned histogram of column *j* of the predicted image and 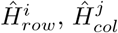 are the same for the ground-truth image, we can approximate the row and column probability density function (PDF) using

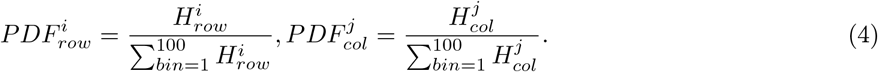

This operation results in *M* + *N* PDFs (one for each row and one for each column). Similarly, *M* + *N* PDFs are calculated for the ground truth image.

Provided with *M* PDFs for the rows from the predicted image and their equivalent row 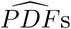 from the ground truth image, we can calculate the KL divergence between *M* such row pairs and *N* such column pairs using

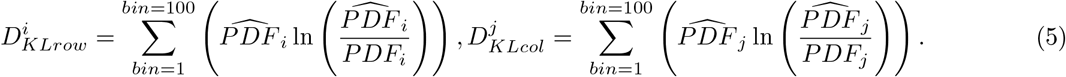

Similar to the weighted L1 loss, we use network driven parameters 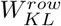 and 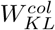, which are 1*× M* and 1*× N* respectively. The attention based weight filtering process is described by

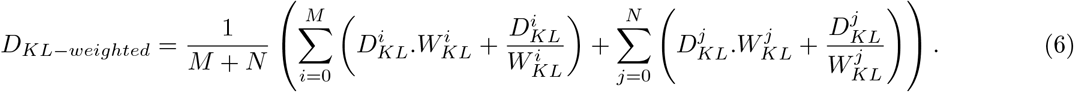

For both L1 and KL losses, the loss is designed such that the weight matrix can only take non-zero values and is scaled between 0 to 1 before multiplication. We then follow [20] to combine the two weighted losses using scaling factors λ_1_ and λ_2_. We found λ_1_ =1 and λ_2_=0.6 empirically provided the best results and learning the scaling factors did not improve the performance. The final loss function can written as:

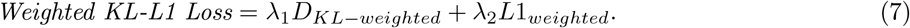

#### 3.3.3 Learnable Weight Matrix

The weight matrices are initialized with random values across all dimensions. The L1 weight matrix is passed through the last DRHT block, which accepts 128×128 sized input and learns to produce the ideal L1 weight map. For the KL weight matrices, we use a stack of 6 linear layers of size 128 for the row weight matrix and a similar set of 6 layers for the columns weight matrix which learn the weights in order to understand rows and columns of importance. The back-propagation occurs as a result of the predicted sinogram as well as the predicted weight matrices. The progression of the weight matrices during training is illustrated in **Figure 7**. We observe that in the learnt weight matrices, L1 weights are higher for areas with subject and lower in the flatter regions. Conversely, the column KL weight matrices have higher weights where there is no subject and lower weights in signal rich regions. The row KL weights, despite having no discernible pattern, helped in improving peak signal to noise ratio (PSNR) marginally. For **Figure 7**, we tested the models after epochs 1,3 and 100 for all 128×128 patches of a given sinogram for all the three datasets. We notice that KL weights are learnt early and minimal change occurs after epoch 30. L1 weights on the other hand, start poorly and then converge to the regions of the subject.

**Figure 7.**
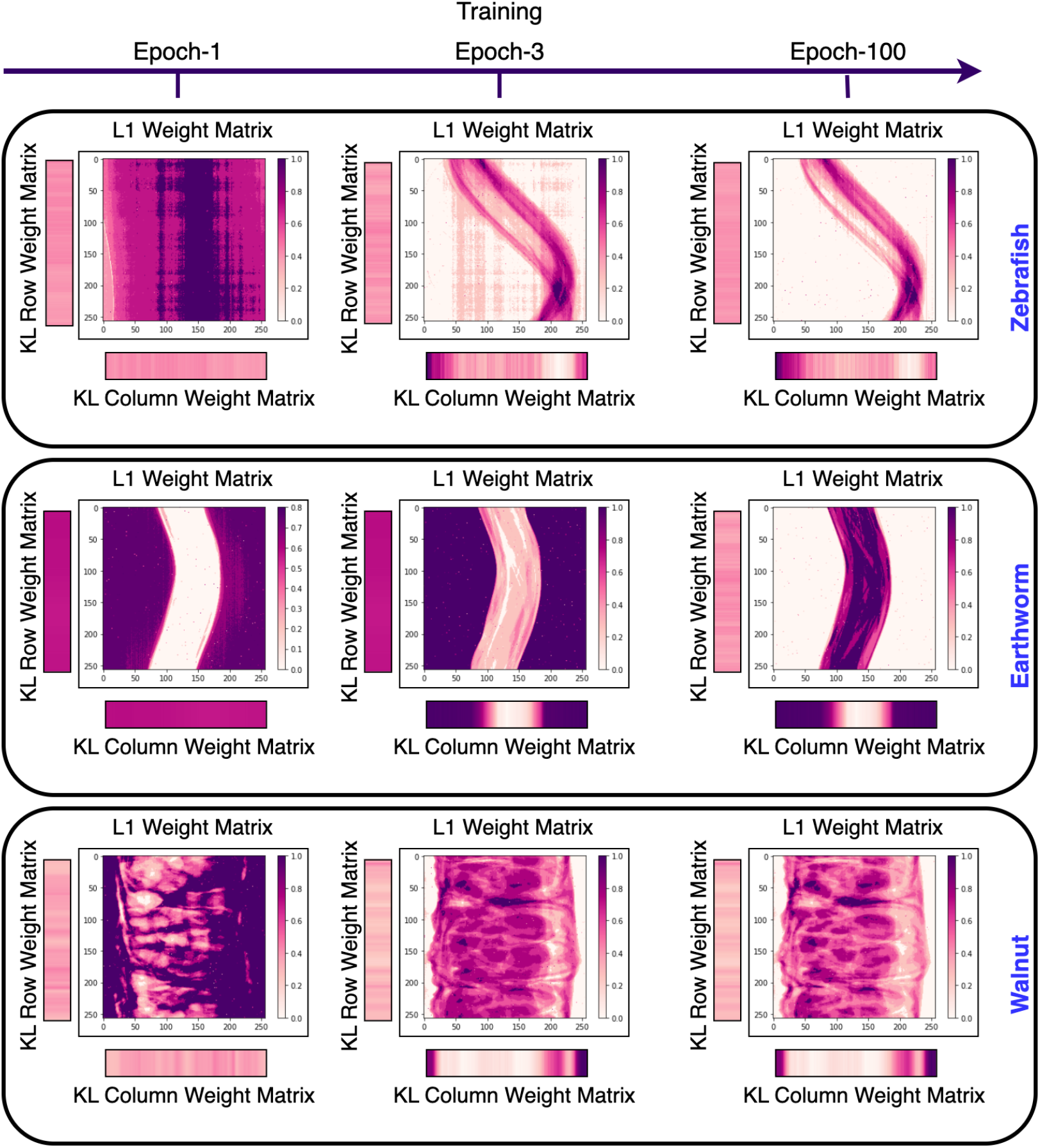
L1 and KL Weight matrices with weights scaled between 0-1 at the end of epoch-1, epoch-3 and epoch-100 for the three datasets showing the learning process. The matrices learn to pay attention to complementary areas and together improve the artifact removal training.

### 3.4 Experiments and Results

We perform two sets of experiments to substantiate our proposed pipeline. First, we conduct a set of ablation studies to determine the need for each of the components in the DRHT network as well as the ablations for the KL-L1 weighted loss module. Then we compare our DRHT against the state of the art models.

#### 3.4.1 Ablation Studies

The three components of DRHT, namely the hierarchical structure (U-shaped), NW-MSA (Transformer blocks), and Dense Residual Blocks and their combinations are tested through multiple models. **Table 2** compares the use of the three components and their respective inference PSNR are averaged over 16.3K Zebrafish samples, 6.1K walnut samples and 8.9K Earthworm samples for 8× upsampling. It is evident that using all three of the components yields the highest PSNR. Additionally, switching the traditional loss function (L1) with KL-L1 resulted in improvement of PSNR scores for both our model and the current state of the art model (Uformer[45]). In the next ablation, we compare weighted KL-L1 against traditional losses and their weighted versions. **Figure 8** illustrates the PSNR gain over the first 102400 iterations by which point all of them had plateaued. While the combination of weighted KL-L1 required more iterations to settle, due to tuning of more weight matrix parameters, it performed the best overall and notably outperformed conventional loss functions like L1 and MSE.

**Table 2:**
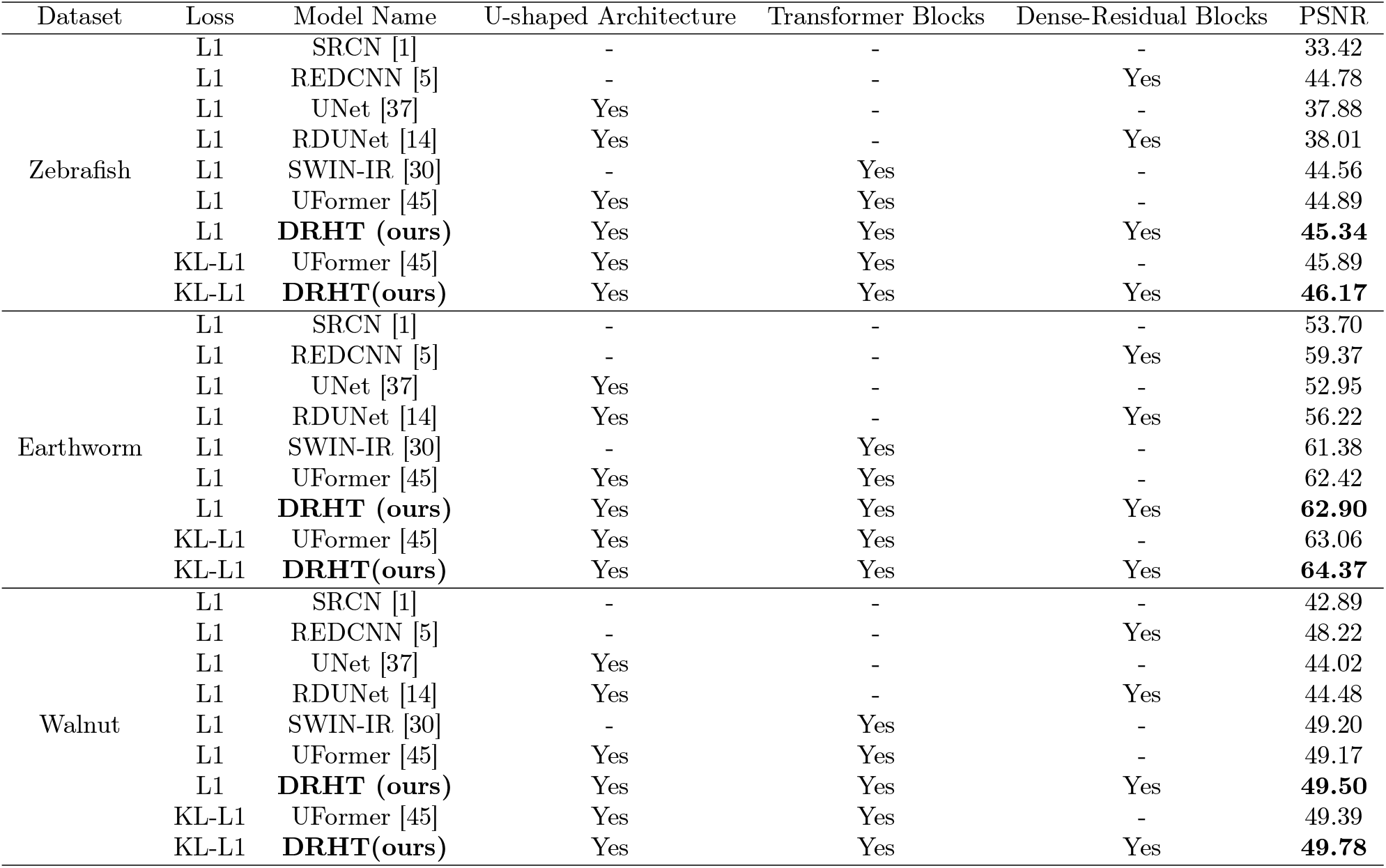
Model Component Ablation for 8× Upsampling of Sinograms

**Figure 8.**
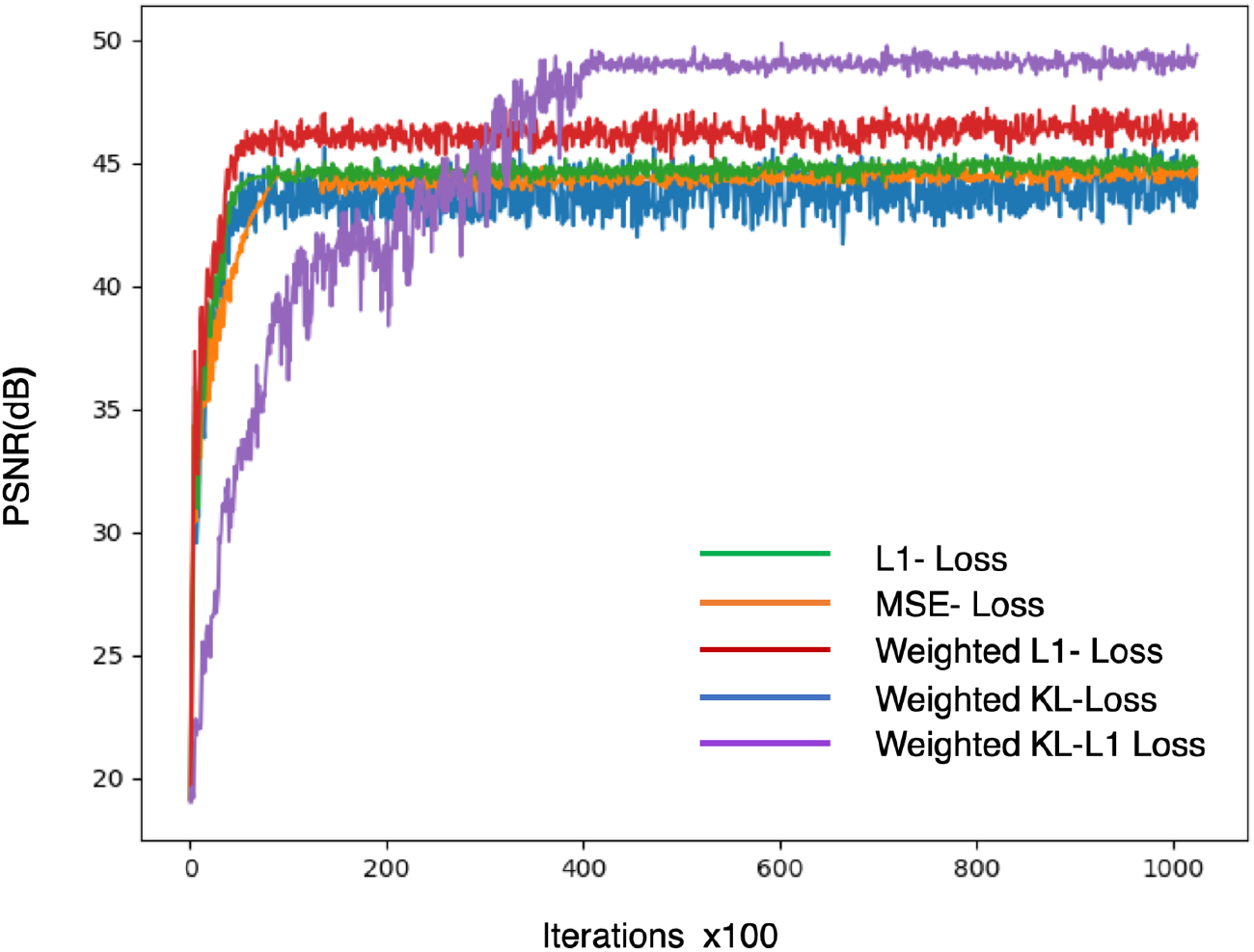
Plots of PSNR gain during training of 2× DRHT with different loss functions for the Zebrafish dataset.

#### 3.4.2 Comparison with State-of-the-art (SOTA) Models

In the second set of experiments, we compare DRHT against popular models like UFormer [45] and REDCNN [5], RDN [53] and a Bicubic baseline using PSNR and structural similarity (SSIM) based metrics. The comparison here is done across 2×, 4× and 8× angular upsampling. The quantitative results of this comparison are presented in **Table 3**. We observe from **Table 3** that DRHT outperforms state of the art models in the sinogram-domain angular upsampling task across multiple scales and datasets. We noticed that that there is an average PSNR increase of 17.73dB and SSIM increase of 0.161 over the Bicubic inputs for the sinogram-domain data.

**Table 3:**
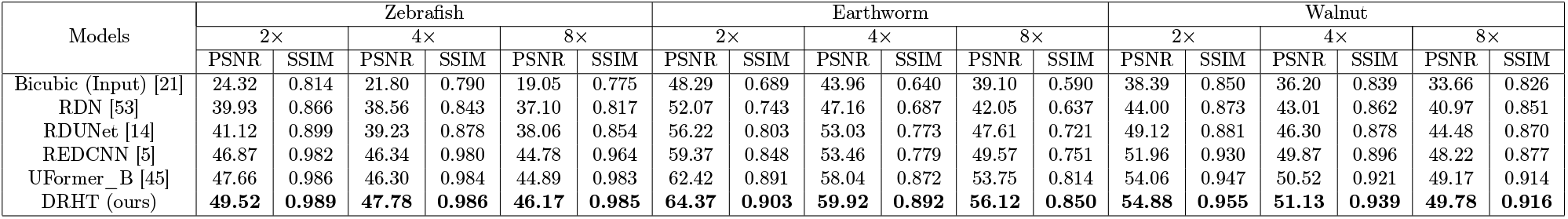
PSNR and SSIM values of test sinograms averaged over 5 runs. **Bold** highlights the best performance

In order to facilitate a fair qualitative comparison, we reconstruct the denoised sinograms and zoom into a specific area of high detail for observing the effect of the models. For the Zebrafish dataset, reconstructions were performed using parallel geometry with the gridrec algorithm in the Tomopy toolbox [15]. Similarly, for the Walnut dataset, we utilized the Astra toolbox [44] to perform a cone-beam geometry based FDK reconstruction and for the Earthworm dataset, we performed the reconstruction using the cone-beam geometry and FDK algorithm in Dragonfly^4^ software with the parameters provided by the authors of the dataset. **Figure 9** (8x),

**Figure 9.**
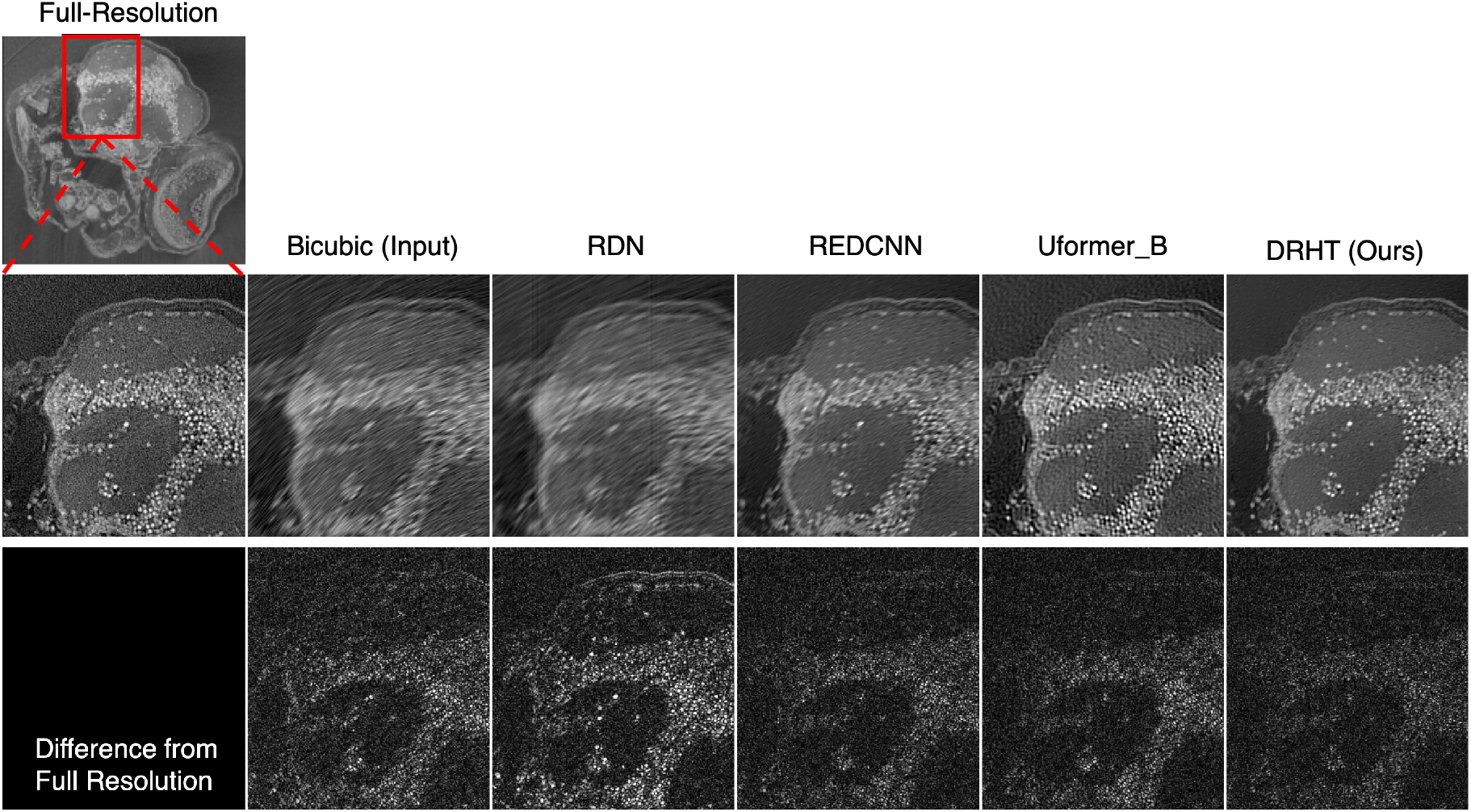
Reconstructed (Transverse) outputs and differences of Zebrafish (slice-700) upsampled from 8× un-dersampled (188 angles) sinogram using various models (Zoom for details).

Figure 10. (4x) and **Figure 11** (4x) illustrate the reconstructions and their residuals provided by the models in comparison. From the qualitative comparison of the reconstructed sinograms, we notice that the DRHT output was the closest to the full-resolution target. Additionally, streaking artifacts were significantly subsided. The quantitative metrics presented in **Table 4** support this and indicate a strong new benchmark for sinogram domain angular upsampling of Micro CT images.

**Figure 10.**
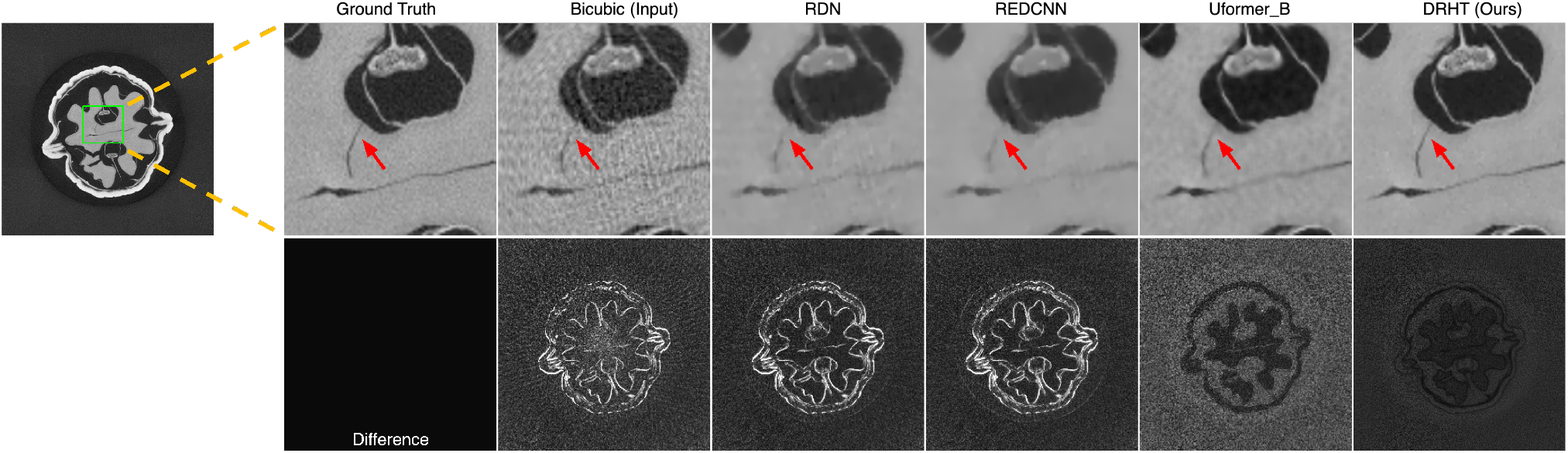
Transverse reconstructions and their residuals from the Walnut dataset (slice-290). The sinograms are 4× upsampled (from 375 angles) before reconstruction. The arrow illustrates the capability of DRHT model to recover fine details with sharpness unlike Uformer (blur), REDCNN and RDN (discontinuity) and Bicubic (artifacts).

**Figure 11.**
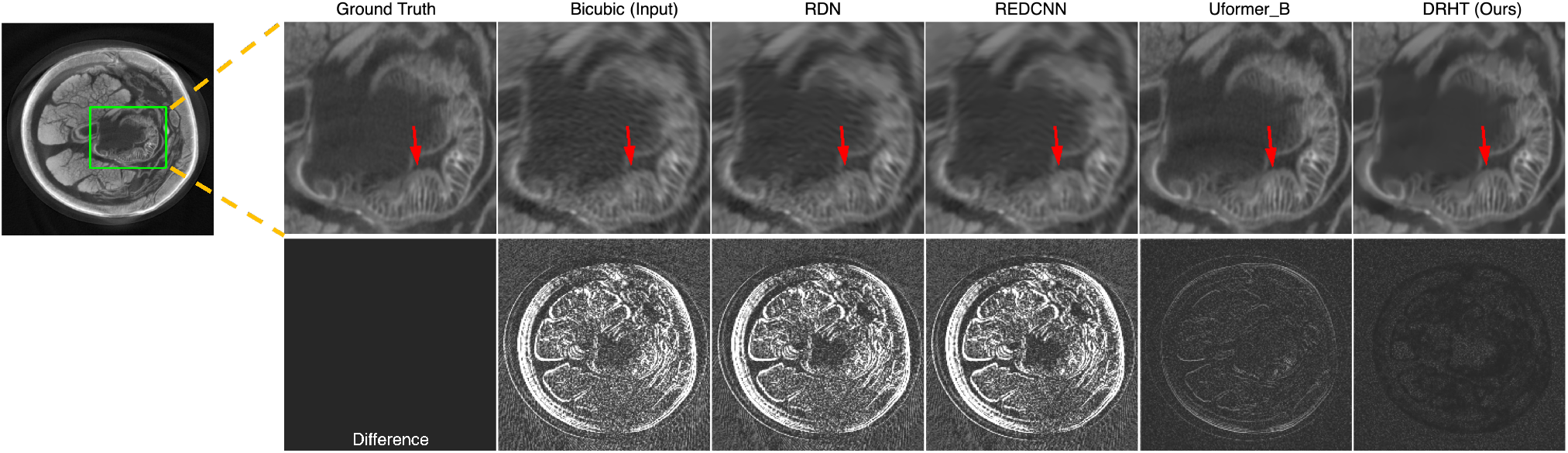
Transverse reconstructions and their residuals from the Earthworm dataset (slice-870). The sinograms are 4× upsampled before reconstruction. The arrow illustrates the capability of DRHT model to separate fine scaled epithelial cell regions in the Earthworm.

**Table 4:**
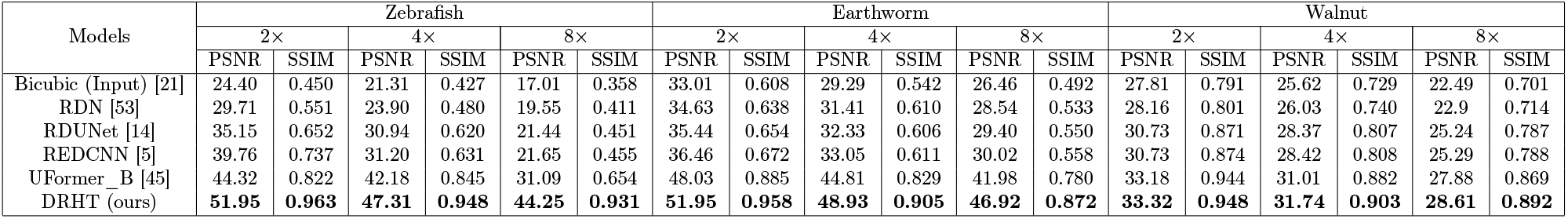
PSNR and SSIM values of reconstructed images obtained from upsampled sinograms averaged over 5 runs. **Bold** highlights the best performance

### 3.5 Observations

The current state of the art architecture, Uformer, performs considerably well on both quantitative and qualitative measures. However, upon closer examination of the reconstructed images, inconsistencies in flat regions become apparent. This can be seen in the residual images of earthworm and walnut datasets. DRHT performs much better in these scenarios and produces images closer to the original full-resolution. In some cases, DRHT-produced images are cleaner than the original ground-truth without elimination of any finer details.

RDN and REDCNN models are not very robust and had varying performances across the datasets. They both performed poorly on the earthworm dataset and had a lot of inconsistencies with edges and sharp features while with the Zebrafish dataset, REDCNN was able to address some of the rotational artifacts.

## 4 Limitations

The use of a deep learning neural network in a supervised pipeline like ours limits the generalizability of the trained model over multiple datasets. For example, we cannot apply the model trained on Zebrafish data to the Walnut or the Earthworm datasets. However, through our experiments, we have demonstrated the capability of our approach when trained on individual datasets at various scales. The training procedure relies on the availability of sinograms which may not be the case for multiple datasets. However, in settings where fast data acquisition is the objective, the DRHT model can be utilized. Additionally, the usage of learnable weight matrices increases the computational requirements due to the calculation of row-wise and column-wise histograms in addition to the KL and L1 losses. This can be observed with the comparison of time taken for iterations presented in **Table 5**. The gain in PSNR comes at higher computational cost but considering the high level details presented by our model in the visualised reconstructions, the requirements are justified.

**Table 5:** Parameters and Time Specifications for Batch Size of 8 on Nvidia GTX 2080-Ti

| Model | Loss | # of Parameters | # of iterations per second |
| --- | --- | --- | --- |
| RedCNN | L1 | 1,848,865 | 7.10 |
| Uformer | L1 | 50,879,216 | 4.45 |
| Uformer | KL-L1 | 57,950,610 | 3.95 |
| DRHT | L1 | 53,419,409 | 3.37 |
| DRHT | KL-L1 | 60,490,803 | 2.82 |

## 5 Conclusion

We have proposed a novel deep-learning model with a U-shaped hierarchical structure for multi-scale feature extraction, non-overlapping window based transformer blocks for identifying long-range feature interactions, and residual dense blocks for spatially local feature extraction and deeper flow of gradients. We applied this model purely in the sinogram domain and empirically showed the significance of each of the sub-units through ablation. Additionally, we further improved the task of micro-CT angular upsampling through the use of a novel noise-aware KL-L1 loss combination that relies on weight matrices for loss calculation.

## Acknowledgement

This work has been supported by the NIH/Office of the Director R24 grant# 5R24OD018559.

## Footnotes

1 https://zenodo.org/record/2686726#.ZC8QJ-zMK8o

2 http://gigadb.org/dataset/100092

3 https://datadryad.org/stash/dataset/doi:10.5061/dryad.4nb12g2

4 https://www.theobjects.com/dragonfly/index.html

